# Harmonizing Clinical Sequencing And Interpretation For The Emerge III Network

**DOI:** 10.1101/457523

**Authors:** The eMERGE Consortium, Richard A. Gibbs, Heidi L. Rehm

## Abstract

**Background:** The eMERGE III Network was tasked with harmonizing genetic testing protocols linking multiple sites and investigators.

**Methods:** DNA capture panels targeting 109 genes and 1551 variants were constructed by two clinical sequencing centers for analysis of 25,000 participant DNA samples collected at 11 sites where samples were linked to patients with electronic health records. Each step from sample collection, data generation, interpretation, reporting, delivery and storage, were developed and validated in CAP/CLIA settings and harmonized across sequencing centers.

**Results:** A compliant and secure network was built and enabled ongoing review and reconciliation of clinical interpretations while maintaining communication and data sharing between investigators. Mechanisms for sustained propagation and growth of the network were established. An interim data freeze representing 15,574 sequenced subjects, informed the assay performance for a range of variant types, the rate of return of results for different phenotypes and the frequency of secondary findings. Practical obstacles for implementation and scaling of clinical and research findings were identified and addressed. The eMERGE protocols and tools established are now available for widespread dissemination.

**Conclusions:** This study established processes for different sequencing sites to harmonize the technical and interpretive aspects of sequencing tests, a critical achievement towards global standardization of genomic testing. The network established experience in the return of results and the rate of secondary findings across diverse biobank populations. Furthermore, the eMERGE network has accomplished integration of structured genomic results into multiple electronic health record systems, setting the stage for clinical decision support to enable genomic medicine.

## INTRODUCTION

The identification, interpretation and return of actionable clinical genetic findings is an increasing focus of precision medicine. There is also growing awareness that the discovery of genes underlying human diseases is dependent upon access to samples from carefully phenotyped individuals with (and without) clinical conditions. As clinical visits provide the ideal opportunity to record patient phenotypes, with appropriate consent, the medical care of specific patient groups can drive the accumulation of clinical data and knowledge of the genetic underpinnings of disease and the penetrance of DNA risk variants. This ‘virtuous cycle’ of data flow from the bench to the bedside and back to the bench will be a key driver of progress in genetic and genomic translation.

While conceptually straightforward, there are many challenges that must be overcome for integrating clinical and research agendas across global populations. Clinical visits are often brief, focused upon measurement related to specific symptoms and constrained by fiscal and practical concerns. On the other hand, ascertainment for research is often open ended, longitudinal, and accompanied by rigorous consent procedures. The types of data that are recorded for each purpose can be different in both depth and quality. As a result, ideal research and clinical records often diverge.

A second group of practical obstacles arises from the heterogeneity of sites and tools used to collect patients’ and participants’ data. On many occasions, even straightforward measurements cannot be meaningfully combined when they are derived from different sites, if they are obtained with different instruments, or from different clinical genetic testing laboratories using different molecular reagents. Thus, some information regarding the method of measurement must accompany the measurement data for harmonization, integration and standardization across populations.

Despite these challenges, the desire to improve medical care by advancing genetic discovery provides incentive for data harmonization. The underlying processes, including participant interaction as well as methods for phenotyping, sequencing, and genetic variant interpretation therefore need to be studied and standardized. Further, the demands of harmonized data flow, storage, and management must be met. Each process must also attend to the tension between respect for patient privacy (e.g. HIPAA laws) and the ability to access data to facilitate research. A list of practical obstacles is presented in Table 1.

**Table 1:**
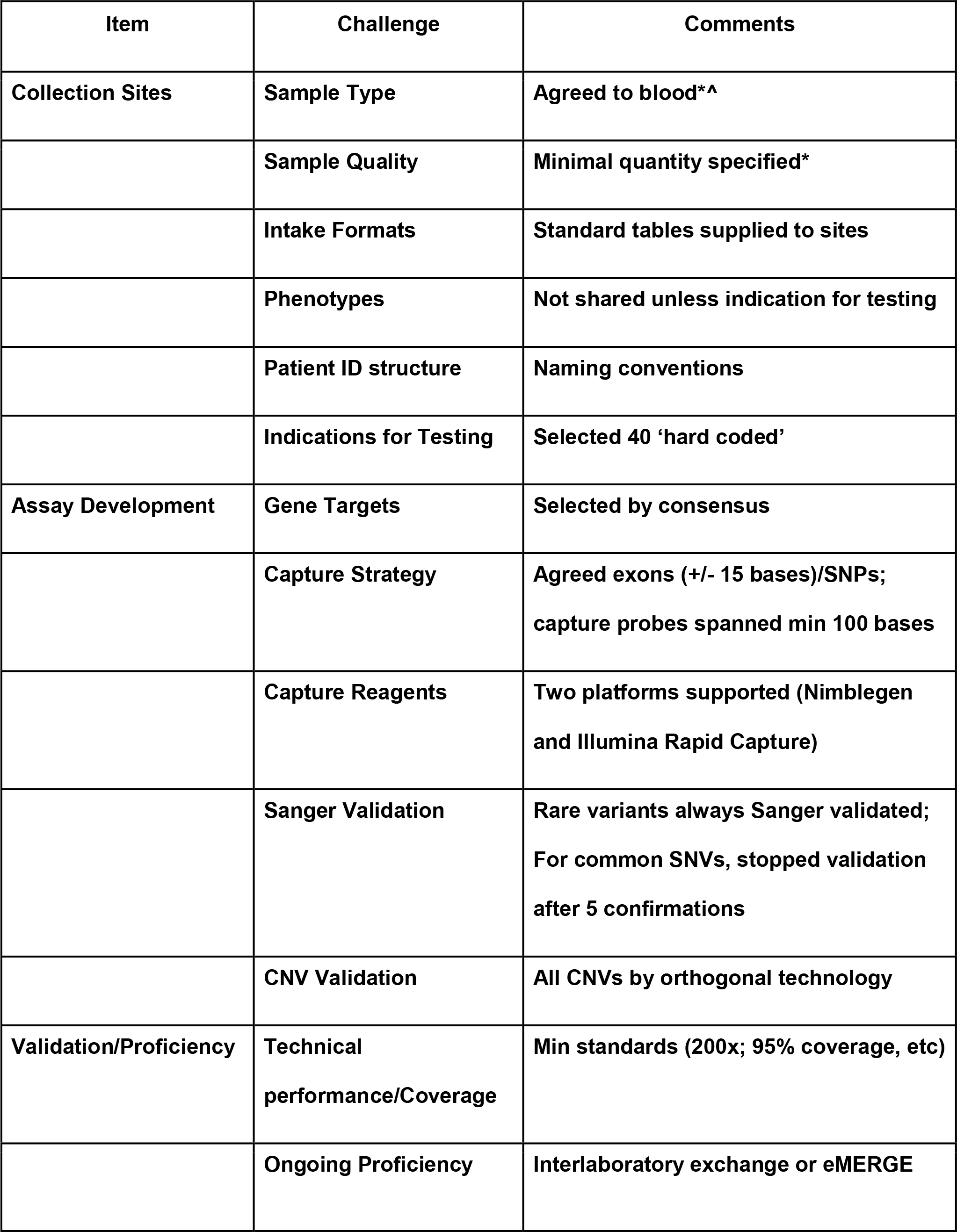
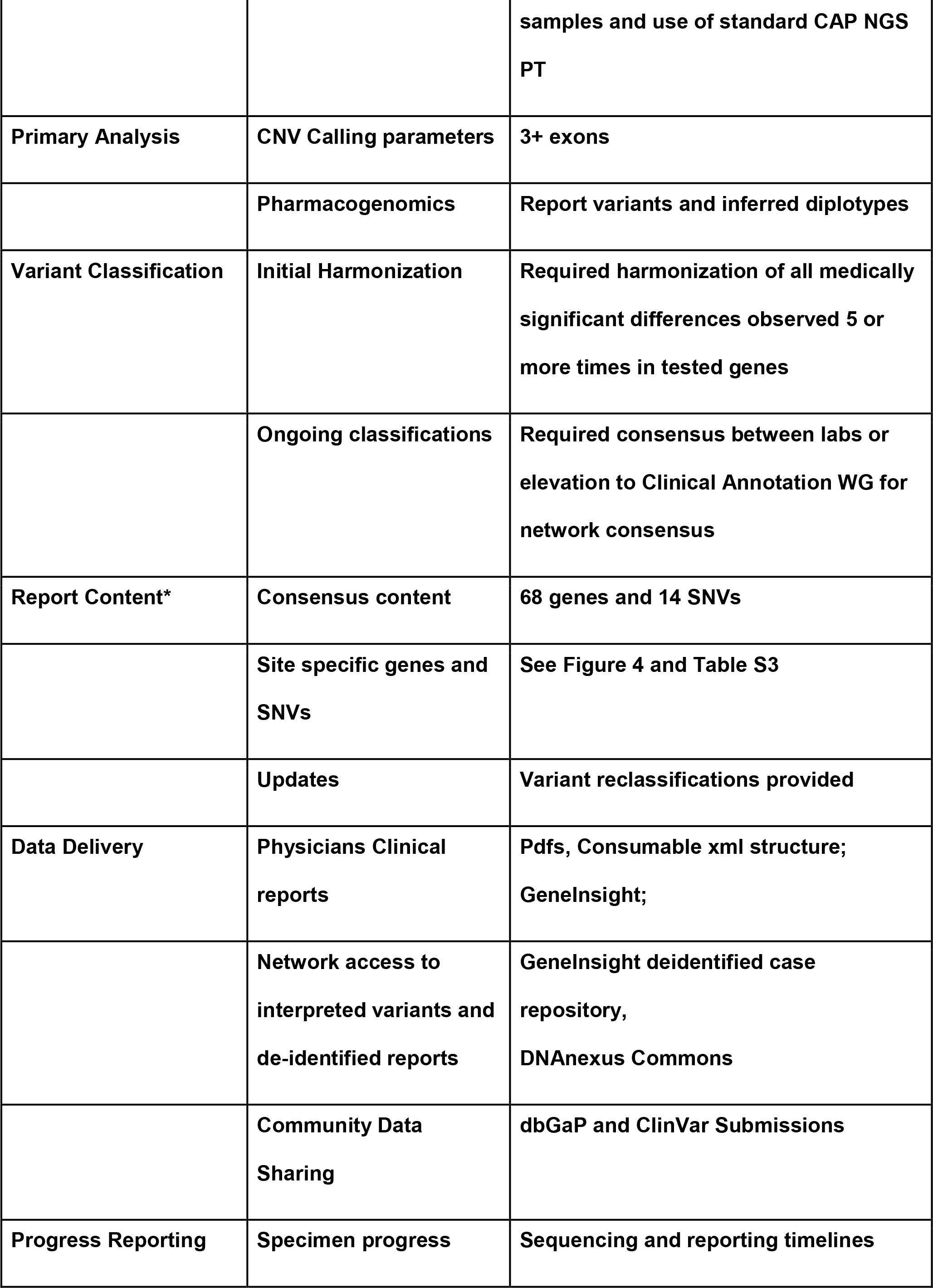
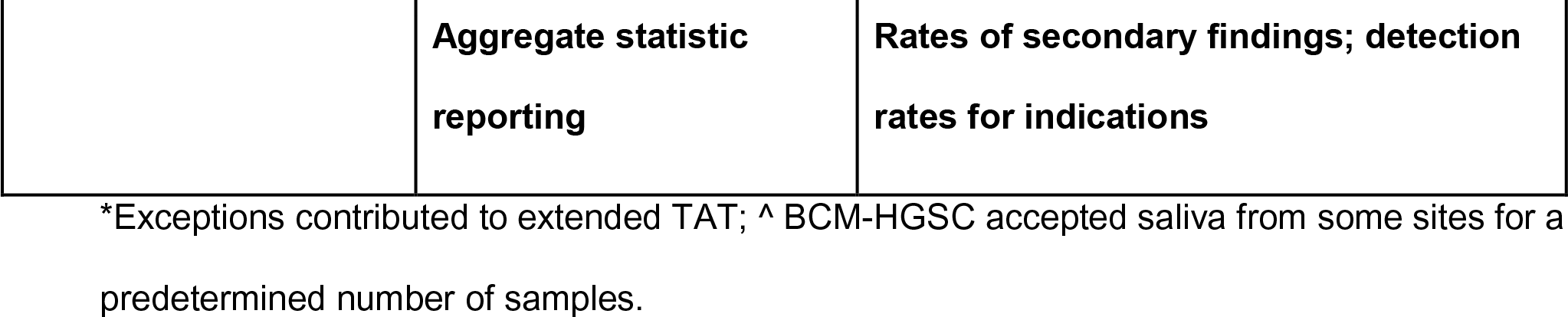
Items to be harmonized.

The current phase (III) of the United States National Institute of Health’s Electronic Medical Records and Genomics (eMERGE) program^1^ aims to study and improve these processes for delivery of clinical and research data, in a multi-center network, while providing actionable genetic results derived from a next-generation sequencing platform to eMERGE research participants. The network builds upon experience with participant consent, obtaining clinical data from the EHR, genotyping and return of results, expanding processes to inform care and catalyze research.

## SUBJECTS AND METHODS

(More details of certain methods are included in Supplementary Material)

### (i) eMERGEseq Panel Overview

#### Panel Design and Content

A gene panel comprising a total of 109 genes and approximately 1,400 SNV sites was informed by network input. The design process considered potential actionability of findings and local research interests, as well as gene size. The 109 genes included 56 based upon the American College of Medical Genetics and Genomics (ACMG) actionable finding list^2^. Additionally, each site nominated 6 genes relevant to their Specific Aims, including discovery-focused genes associated with clinical phenotypes in need of further study. All nominated genes apart from titin *(TTN),* which was excluded due to its size, were included in the final panel design for a total of 109 genes. Further, eMERGEseq content included several categories of single nucleotide variants (SNVs): 1) ancestry informative markers and QC/fingerprinting loci (N=425), 2) a suite of SNVs selected to inform HLA type (N=272), 3) pathogenic SNVs in genes not included on the panel for which return of results was planned (N=14), 4) pathogenic or putatively pathogenic SNVs in genes not included on the panel for which return of results was not planned (N=55; for some, penetrance is poorly understood), 5) SNVs related to site-specific discovery efforts (N=718), and 6) pharmacogenomic variants (N=125), selected based on potential actionability, allele frequency and space available on the platform. A summary of all eMERGEseq content can be found in Table 2, with additional details provided in Table S1. All sequence and SNV data are shared across the network for research, and a subset of the content, namely the clinically actionable variants associated with disease or drug response, are included in clinical reports for return to the participants.

**Table 2:**
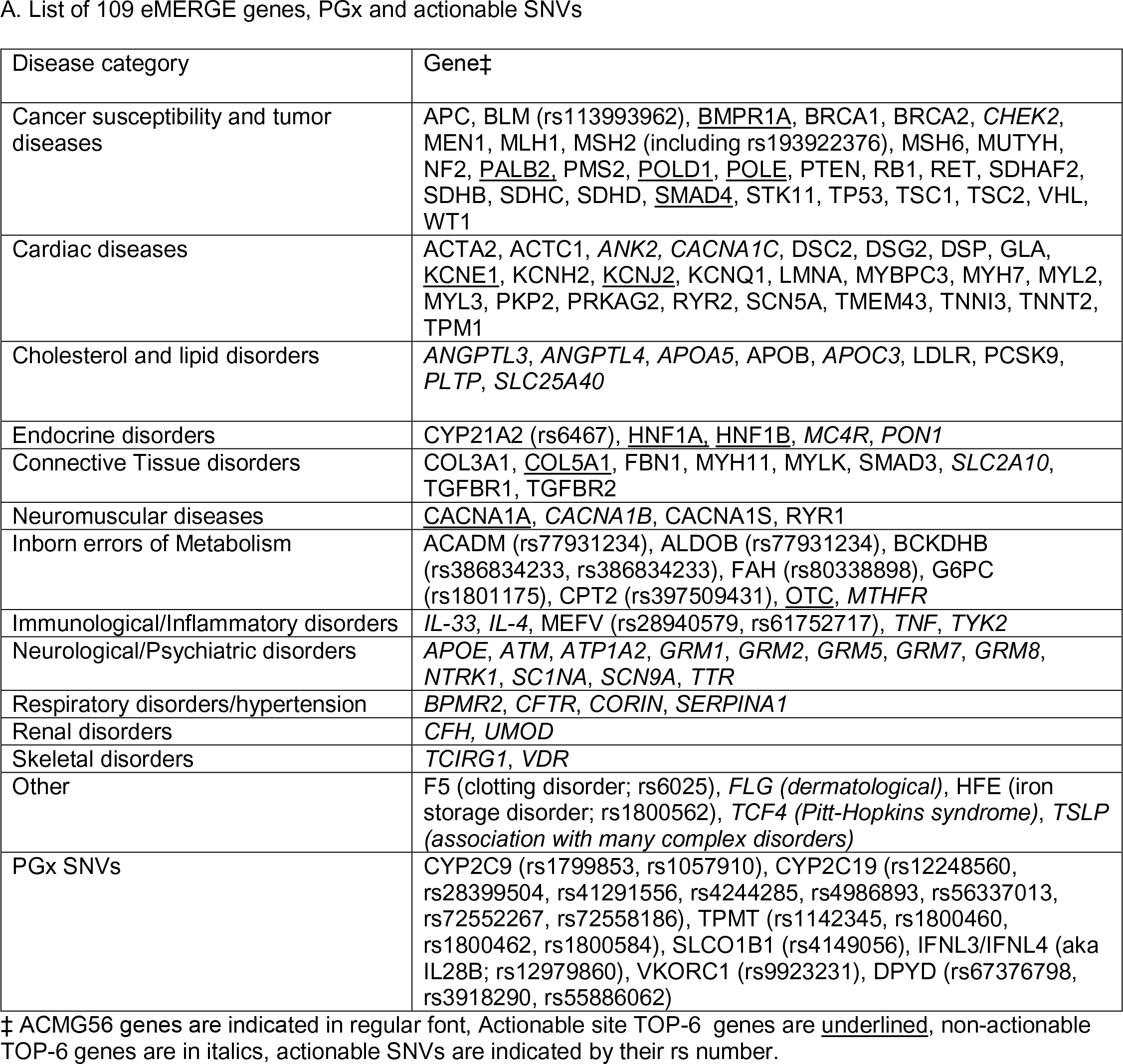
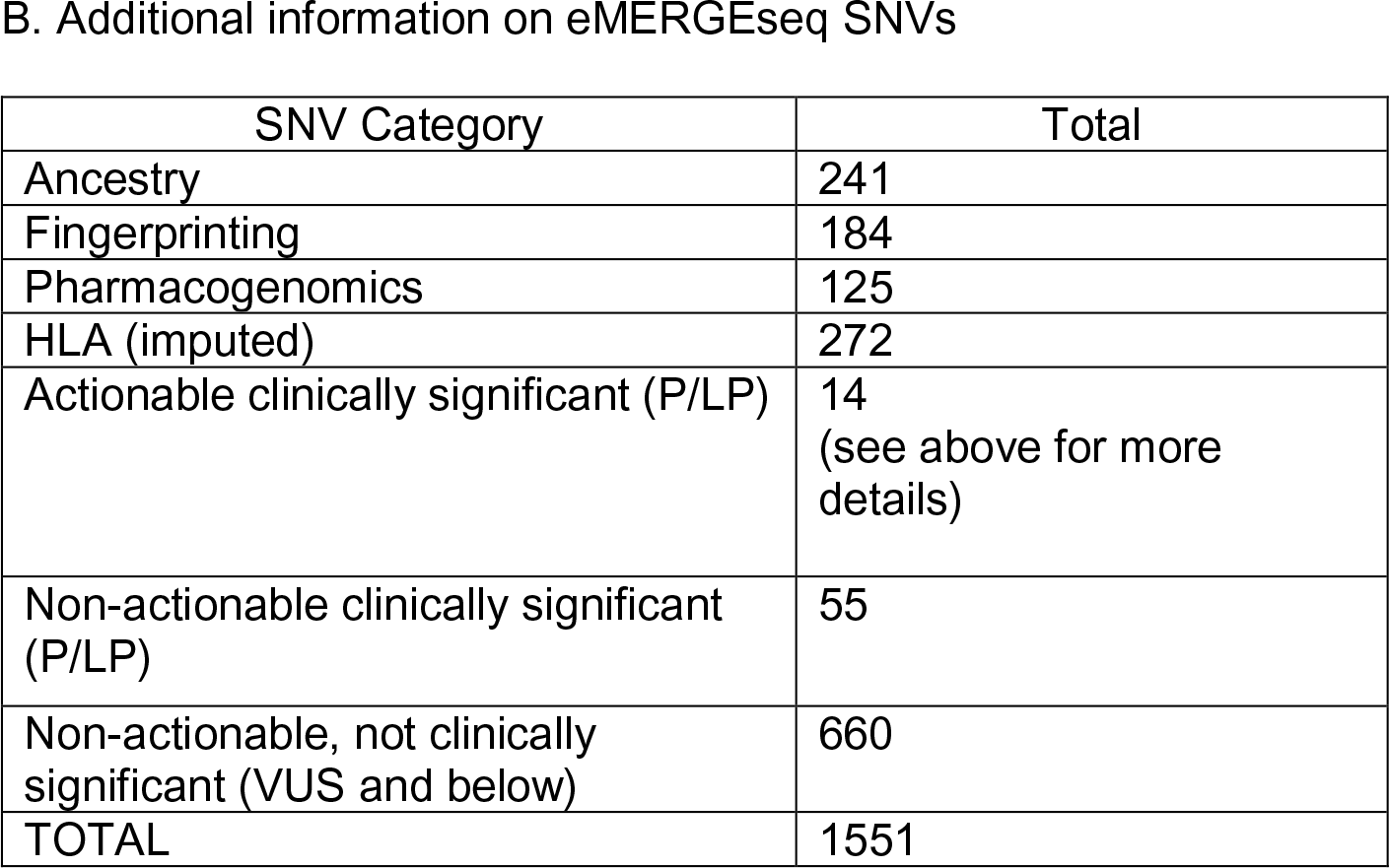
Genes and 914 SNVs on the eMERGEseq Panel.

### (ii) Panel Sequencing

#### Reagent

The gene and SNV list was used to direct construction of targeted capture platforms at two sequencing centers (SCs): The Baylor College of Medicine Human Genome Sequencing Center [BCM-HGSC], Houston TX; Broad Institute and Partners Laboratory for Molecular Medicine, Cambridge, MA. Broad used Illumina Rapid Capture probes for this panel and the BCM-HGSC used Roche-Nimblegen methods. Each group created in-solution capture probes spanning the entire targeted regions of the eMERGEseq panel. Probes were designed to be complementary to specified exons or SNV sites with a minimum span of 100 nucleotides. Tiling was limited to exonic sequence and analyses included +/− 15 intronic flanking bases (Figure S1).

#### Sample preparation

Clinical sites were requested to submit 2ug of extracted DNA within a concentration range of 30-50 ng/ul. Although DNA derived from blood was the specified sample for the program, BCM-HGSC revalidated the clinical assay and accepted saliva as a DNA source for a limited number of cases due to clinical site requirements. Once received by the sequencing center, specimens were quantified using a picogreen assay, and quality was assessed by gel. Specimens with a minimum of 600 ng of DNA that did not display high levels of degradation passed sample QC and were accepted for eMERGEseq testing.

#### Sequencing and Primary Analysis

Samples from DNA capture using the custom capture reagents were sequenced using standard Illumina technologies. Post-sequence processing at each site utilized preferred alignment and variant calling algorithms. The variant calling pipeline at Broad incorporates Picard deduplication, BWA alignment, and GATK variant calling for SNVs and short InDels^3^. At the BCM-HGSC the alignment using BWA-MEM and variant calling using Atlas were instantiated within the Mercury Pipeline^4^.

#### Panel Fill-in

A common set of reference samples were initially sequenced at each SC. The chosen parameters to monitor performance were coverage of targeted sequence and percentage of the targeted bases at or above 20X coverage. Both groups sequenced cohorts of control samples and identified systematically poorly-covered bases as those with less than 20X coverage in >10% of tested samples. Based on this conservative threshold, both groups went through a process of enriching with more targeting probes (‘fill-in’), to boost underperforming regions, prior to final validation. The reagent performance is described in Table 3, with additional details in supplementary Table S2.

**Table 3:** Assay performance and optimization at the sequencing sites.

|  | BCM-HGSC |  |  | Broad |  |  |
| --- | --- | --- | --- | --- | --- | --- |
|  | Acceptance<br>Criteria | Original | Low Input | Acceptanc<br>e Criteria | Measured<br>at ~250X<br>MTC | Measured<br>at ~400X<br>MTC |
| Assay sensitivity (SNV + indel) |  | 100% | 100% | ≥95% | 100% | 100% |
| Assay sensitivity- CNV |  | 97.7% | 98.3% | n/a | 100%* | n/a |
| Assay specificity (point variant +<br>indel) |  | 100% | 100% | ≥95% | 100% | 100% |
| Assay reproducibility | ≥95% | >98% | >97% | ≥95% | 98.5% | 99.6% |
| % of >20X coverage for targeted<br>regions | ≥99% | >99% | >99% | ≥95% | 99% | 99% |
| Depth of mean coverage | >200X | >200X | >200X | n/a | ≥250X | ≥400X |

\*CNV sensitivity at Broad/LMM is for events ≥3 consecutive exons.

#### Copy Number Variant (CNV) Calling

CNV calling at Partners/Broad was performed using VisCap, which infers copy number changes from targeted sequence data by comparing the fractional coverage of each exon in a gene to the median of these values across all samples in a given sequencing run^5^. BCM-HGSC CNV calls were made via Atlas-CNV, in-house software that combines outputs from XHMM^6,7^ and the GATK DepthOfCoverage tool^6^. Like VisCap, Atlas-CNV infers the presence of CNVs from normalized coverage differences to other samples in the same sequencing batch, and refines these predictions with a pair of quality control metrics^8^. CNV calls were confirmed by orthogonal technology - Droplet Digital PCR (Bio-Rad, Hercules, CA) at Partners/Broad and Multiplex Ligation-dependent Probe Amplification (MRC-Holland, Amsterdam, Netherlands) at the BCM-HGSC. Detected CNVs were filtered based on clinical site’s gene reporting preferences and ClinGen haplosensitivity and tri-sensitivity scores (https://search.clinicalgenome.org/kb/gene-dosage), and then manually reviewed. Partners/Broad required a minimum of three contiguous exons for reporting, BCM-HGSC required two.

#### Analytical Validation

To validate sensitivity, specificity, and reproducibility of the eMERGEseq panel, the performance of both SCs was compared using a common reference sample (NA12878). In addition, each group separately examined previously tested clinical samples, containing known pathogenic variants that were uniquely available to their laboratory. Subsequent additional validation analyses were performed to accommodate lower DNA input amounts, based on sample availability (BCM-HGSC).

#### Ongoing Proficiency

Ongoing proficiency testing involved interlaboratory exchange of previously tested eMERGE samples and CAP proficiency testing for general sequencing platforms with all results concordant to date (see Supplementary Materials: Supplemental Methods).

### (iii) Variant Interpretation

#### General Approach to Interpretation

Variant classifications from both laboratories were based on ACMG/Association of Medical Pathology (ACMG/AMP) criteria with ClinGen Sequence Variant Interpretation

Working Group modifications as well as additional specifications for some of the eMERGEseq genes as established by ClinGen Expert Panels^9^. Additional local data accrued from previous case studies was combined with manual literature and public data review for final decisions. Non-ACMG 56 genes underwent an in-depth clinical curation effort using the ClinGen framework for gene-disease validity assessment^10^.

#### Legacy Variant Interpretation

In order to harmonize prior interpretations and to assess likely ongoing differences, the BCM-HGSC and Partners LMM exchanged data from 1,047 previously interpreted variants in the 109 eMERGE genes and evaluated discrepancies (see results).

#### Ongoing Harmonization

Monthly data exchanges identified any differences of interpretation of non-PGx variants intended for clinical reporting. These discrepancies were reviewed during a bi-weekly interpretation/harmonization teleconference call. Cases of unresolvable variants were presented to the eMERGE Clinical Annotation WG to attempt resolution and/or track their occurrence. All reported variants are submitted to ClinVar with their interpretations.

#### Pharmacogenomics (PGx)

The SCs worked with the eMERGE PGx working group to: select variants to be included on the clinical reports provided to participants; interpret diplotypes; and select drugs for therapeutic recommendations, guided by CPIC guidelines. Twenty PGx variants in seven genes were deemed to be clinically actionable and selected for return to participants. Table S4 includes details of the PGx genes and variants reported and the drugs associated. For two PGx genes, *CYP3A5* and *SLCO1B1,* the gene panel included only one of three variants discussed in the CPIC guidelines. *CYP3A5* was deemed not reportable, as two SNVs important for predicting phenotype for African Americans and Latinos are not included on the gene panel. *SLCO1B1* was deemed reportable, as the one SNV included in the panel serves as a tag SNV for the remaining two SNVs.

The BCM-HGSC included PGx results on individual patient reports, while Partners LMM produced a batch report that accommodates one to hundreds of patients for bulk consumption and EHR integration by sites. Sample PGx reporting formats can be found in the Supplemental Material (Sample eMERGE report HGSC-CL, LMM Sample PGx Batch Report). The CPIC drugs that were included in the PGx report were largely the same with some minor differences (see Table S4).

### (iv) Data Management

#### Sample Intake

Each site was provided barcoded tubes by the SC for DNA shipping. Sample identifiers and metadata were uploaded using an ‘eMERGE requisitioning sheet’ via secure portals^11,12^. The requisitioning spreadsheet contains fields for sample information (name [optional], sex, date of birth/age, US state of residence, site-specific ID), as well as eMERGE-specific metadata including; patient ‘disease area’ (from a list defined by the network - see supplementary material for details), disease status and test indication, eMERGE project ID and barcode number on the tube. An additional option was to add phenotype terms in a free-text field, primarily based on the MonDO ontology and occasionally additional local codes largely derived from Human Phenotype Ontology (HPO) terms (See Supplemental Material: preferred indication terms). A simple .csv file structure was used by both SCs so that sites could upload all metadata at the time of sample batch shipment. For the BCM-HGSC SC, the sample accession was directly into a cloud environment, managed by DNAnexus, while for the Partners-Broad site a custom portal operating in the Broad’s local environment was employed for intake followed by transfer to the GeneInsight system for analysis and reporting; all systems were HIPAA compliant. Local identifiers were then generated to track the samples as they progressed through DNA sequencing and variant calling. Orders were reviewed and approved by the SCs prior to sample shipping and accession. Upon receipt, the samples were subjected to volume and concentration quality control checks.

#### Data Delivery and Reporting

Each SC developed custom reporting methods (see Supplementary Material for examples). Partners/Broad site users have a unique, password-protected account and are only able to view orders and metadata from their own site. The Broad portal authorization procedures are customized to allow for secure transfer of sequencing output files and metadata to both Partners and DNAnexus via APIs. The BCM-HGSC sites are delivered reports from the DNAnexus environment via DNAnexus APIs. Users were provided individual logins for accessing pdf reports and structured content in a harmonized .xml format.

#### GeneInsight

Partners/Broad sites used the commercial tool, GeneInsight (Sunquest Information Systems, Tucson, AZ), for local report management^13^. This tool was configured to create a De-identified Case Repository (DCR) which contains a de-identified record of all cases and associated variants from both Partners/Broad and the BCM-HGSC supported sites.

#### DNAnexus Data Commons

The BCM-HGSC clinical sites were provided with two data access points in the DNAnexus infrastructure One provides a restricted space for accessing PHI-containing clinical reports, while another acts as a general space for the de-identified records of each case and associated variants. Users were provided individual logins and selectively granted access to one or both access points. Data for sites that were served by the BCM-HGSC were provided both .xml and .pdf formats, at the time of reporting. De-identified, structured versions of the Partners-Broad reports are downloaded from the DCR and also stored in the DNAnexus Data Commons projects, creating a comprehensive repository of de-identified clinical reports.

#### Variant Updates

Two complementary mechanisms were developed to enable delivery of variant updates from the SCs to the sites as new evidence leading to a classification change becomes available. At Partners/Broad, individual participant results are stored in an eMERGE-specific instance of the GeneInsight database that is linked to Partners LMM’s GeneInsight instance enabling communication of variant updates^14^. If Partners updates a variant, sites that have signed up receive proactive notification emails if a reported variant identified in one or more of their cases is updated. Hyperlinks are provided in those emails that allow sites to directly access updated information on the variant in each case, which facilitates the choice to return an updated result to a participant. In addition, Partners is generating an .xml file for each variant interpretation change alert, which sites can consume through other electronic interfaces. At the BCM-HGSC, participant results are stored in a database that is routinely queried for variants with new actionable interpretations. If such a variant is found in a previously-reported sample, an amended report is issued via DNAnexus and sites are notified. Variant updates are included in the ongoing variant interpretation harmonization process described above.

### (V) Data Freeze and Raw Data Storage

In order to analyze preliminary results from the eMERGE III eMERGEseq data, an interim freeze of samples sequenced by November 2017 was generated. These 15,754 samples (9633 from Baylor and 6121 Broad-Partners) and the available associated data are described in detail in the Supplementary Material. The associated BAM, xml and vcf files are available on the eMERGE Commons, accessible to sites as well as outside investigators who apply for access (https://emerge.mc.vanderbilt.edu/collaborate/). Data are also submitted to dbGaP for controlled public access.

## RESULTS

### (i) Network Overview

The eMERGE III network established a Clinical and Discovery Platform that consists of 11 clinical study sites, two DNA SCs and a coordinating center (CC) (Figure 1). Participants were enrolled at each site, blood collected, DNA extracted locally and sent to one of two SCs for targeted sequencing. Analysis and interpretation of the DNA sequence data was performed at each SC, and the data returned to the clinical sites for return to participants. Raw data were accrued for data mining purposes by eMERGE investigators and approved affiliates. Subsequently raw data are released to dbGaP and interpreted variants to ClinVar.

**Figure 1:**
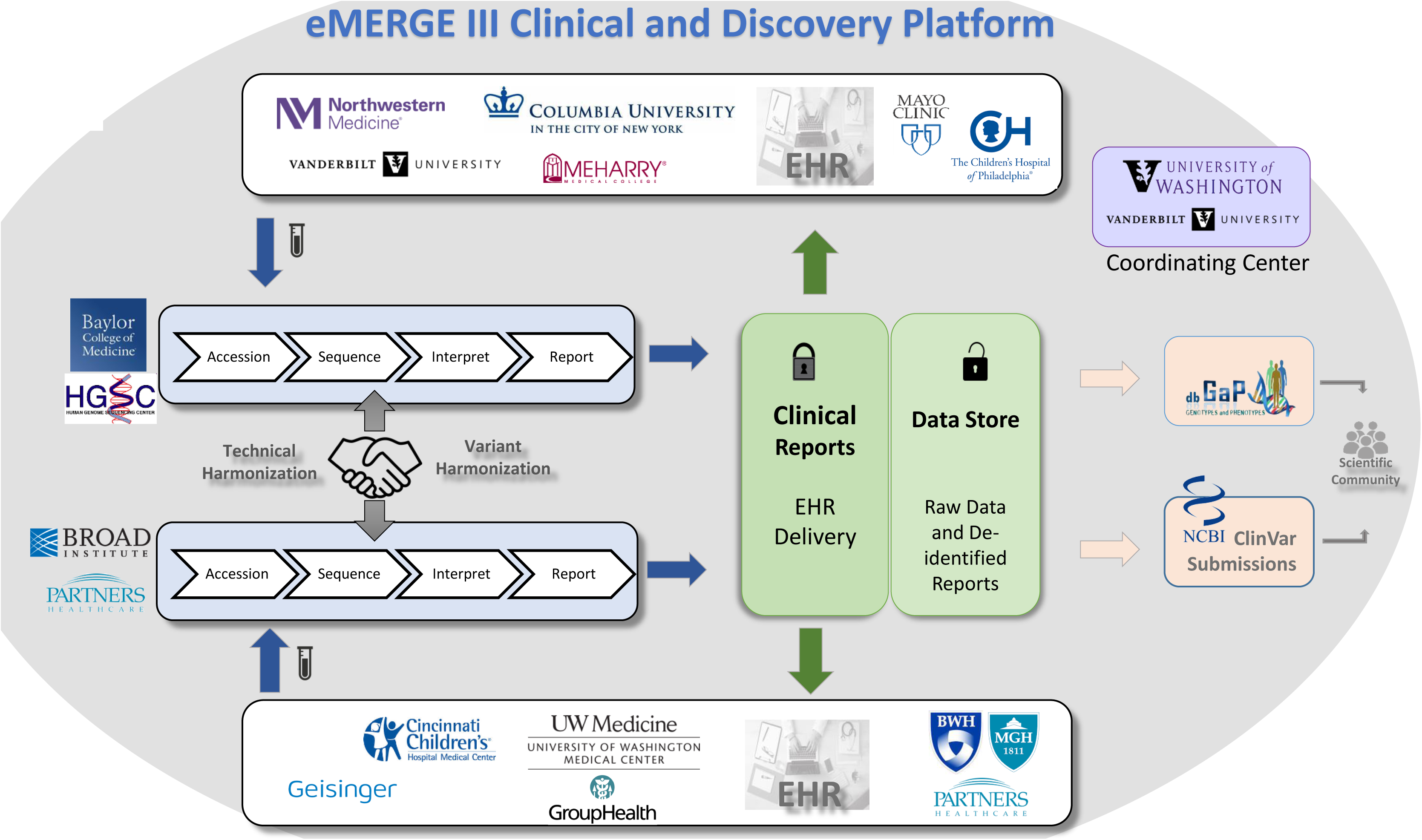
eMERGE III Network Overview. The eMERGE III network is comprised of 11 study sites, two sequencing centers (SCs) and a coordinating center (CC). The different components and processes involved in the data flow across both the clinical and research discovery arms of the network are highlighted in this figure and described in more detail in the network overview section.

An early decision of the program was to utilize DNA capture ‘panels’ of approximately 500 kb, in order to generate genomic data from the eMERGE participants, as an alternative to genotyping, whole exome sequencing (WES) or whole genome sequencing (WGS). This choice reflected a balance between available fiscal resources and a reasonable selection of content to explore return of actionable results and focused discovery efforts. The use of the panel enabled testing of 109 genes and approximately 1400 additional sites of single nucleotide variation in each sample. Across the network, ∽25,000 samples are being assayed, ∽2500 from each site. The study is therefore large enough to allow robust analysis of specific phenotypes, as well as to gain experience with a sufficient number of patients at each site to develop processes to support the return of actionable genetic results.

Prior population studies suggested that the genes included on the panels would reveal thousands of newly identified single nucleotide and structural variants. A small subset of these would be expected to be pathogenic, and the program aimed to report to participants only those variants that were pathogenic or likely pathogenic according to the ACMG/AMP guidelines^15^, or those with actionable pharmacogenomic associations. In addition, it was aimed to provide data from the panel that informed possible pharmacological responses. Each site would have the option of a customized clinical reporting framework, as well as full access to all network data to guide decisions and harmonize interpretations.

This elaborate network reflects a real-world situation, where a full complement of testing, reporting, and research require coordination and harmonization of many components. First, the selection of gene targets and the rules for reporting must agree. Next, the technical aspects of DNA capture and sequencing required standardization and ongoing comparison. The DNA changes must be interpreted and reported with the same conclusions, regardless of where testing occurred. Finally, file structure standardizations and data management practices must be organized. A detailed list of components (Table 1) that require coordination and harmonization illustrates the magnitude of the challenge.

### (ii) Technical Validation of Capture Panels

Coordination and harmonization of the DNA capture panel process at the two CAP/CLIA certified DNA sequencing laboratories was demanding because in addition to different DNA capture reagents, the local processes of sample preparation, library construction, hybrid capture, and sequencing represented complex workflows with many variables. As an alternative to compelling each laboratory to adopt unfamiliar methods, the harmonization was achieved through phases of coordinated design, comparing initial high level technical performance and via ongoing monitoring of proficiency (Figure 2a). The harmonization process aimed to reduce any impact on the overall program of the heterogeneity of capture reagents or sequencing methods between two sites and for the end users to be able to compare data from each laboratory without batch effects.

**Figure 2:**
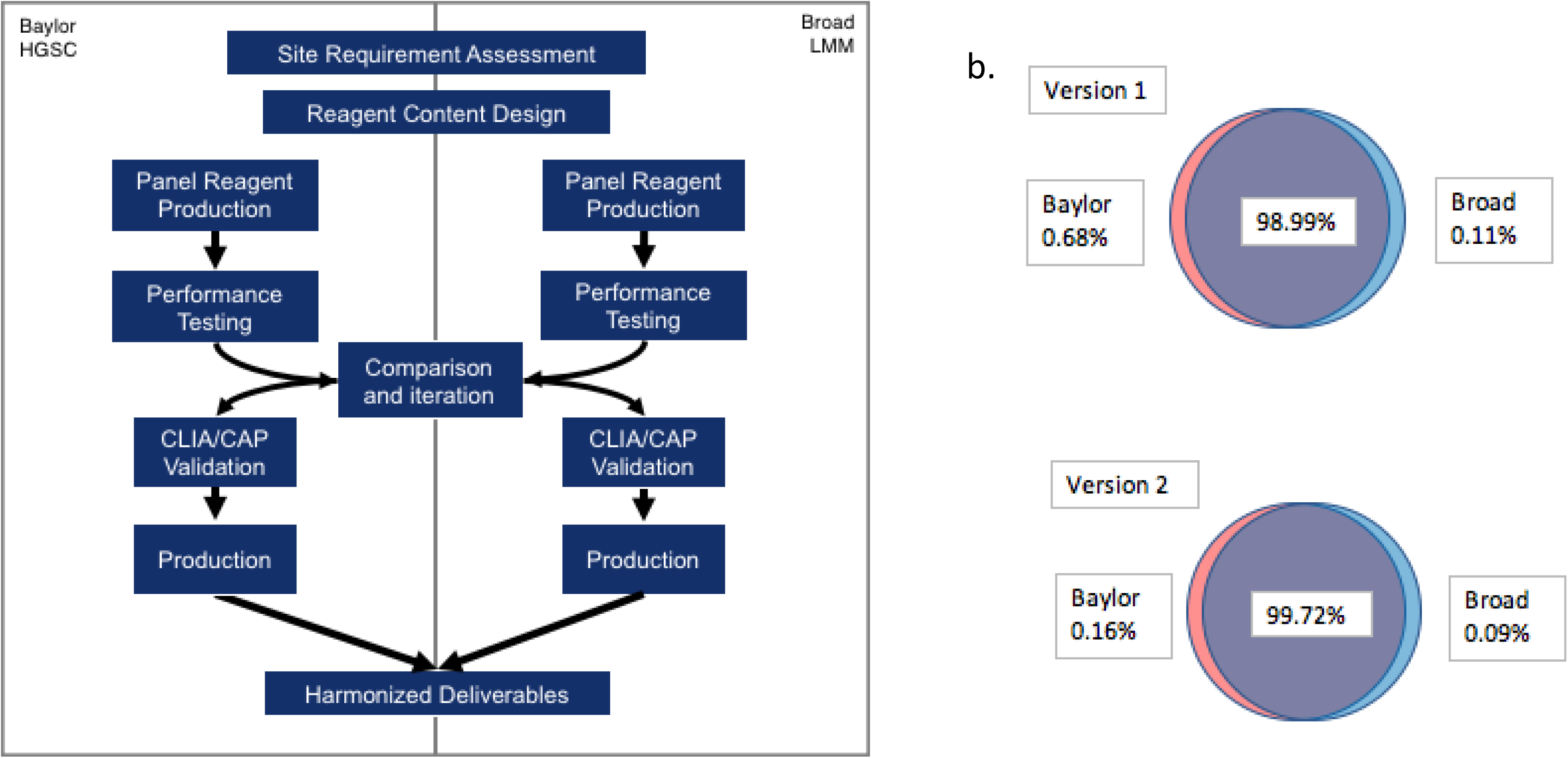
eMERGEseq Panel Test Development and Validation. a. Technical Harmonization of Two DNA Capture Panels. Coordination and harmonization of all the components of the DNA gene capture panel process at the two sequencing centers. b. Base Coverage. Percentage of bps covered ≥20X Across Sequencing Centers. % of bases in the panel targeted region covered in each version of the panel design and the extent to which these bases overlap between the genome centers is shown. Version 2 is the final version used for data generation.

Design was coordinated by first agreeing on the intended limits to reporting, e.g. number of bases adjacent to exons to be reported (see methods, Figure S1). Each laboratory employed slightly different criteria for the selection of the range of transcripts to be tested, reflecting a lack of harmony of public databases. Possible differences in design were resolved by selection of the union of all possible exons to be considered, and validated by iterative sharing of the capture design files (‘bed files’). The detailed design specifications can be found in Table S1.

Preliminary testing of the technical performance of the two capture reagents utilized both local test samples and a shared sample reference set (see methods). The technical performance was shared between the SCs by measuring the coverage of individual bases and other key technical metrics (Table 3, Table S2). Overall sequence coverage goals and the extent to which poorly covered regions could be tolerated were agreed upon *a priori,* and the technical comparison was straightforward between SCs. In general, the sequencing reagents performed well, although the presence of some uncovered bases in the first panel designs led each group to modify the initial reagents to optimize performance (Figure 2b). Throughout, the comparative performance of the two reagents informed the progress of technical development and illustrated the synergism from closely monitoring similar processes.

For final validation, both groups measured overall sensitivity and specificity on a reference sample (NA12878) as well as sensitivity to detect known pathogenic variants from previously tested clinical samples that were uniquely available to them. Groups also incorporated evaluation of variance in processing including varying coverage from ∽250X to 400X (Broad) and input amounts of 250 ng and 500 ng (Baylor). Summary results of the respective validation studies are shown in Table 3. Panel optimization results and coverage analyses can be found in T able S2. The impact of the ∽0.2% of targeted bases that were not effectively covered via the optimized panel designs was evaluated by the network for impact on clinical decision making. The majority of missing data was judged to be of little consequence although small regions of some genes (e.g. *RYR1, CACNA1B*) could not be recovered by either platform.

Once the data production phase of the program was initiated, the ongoing performance was monitored by sharing production metrics and via the ongoing CAP/CLIA proficiency program that included exchange of samples and comparison of DNA variation data. As of this publication, mean coverage of Broad production samples is ∽420X, % of targeted bases covered ≥20X is 99.7%, and % of targeted bases with zero coverage is 0.17%. These metrics, collected from >7000 production samples, closely match the performance of the validation set. Mean coverage of the BCM-HGSC production samples is ∽340X, % of targeted bases covered ≥20X is 99.8%, and % of targeted bases with zero coverage is 0.04%. These metrics, collected from >9600 production samples, also closely match the performance of the validation set.

### (iii) Clinical Content Validation and Site-Specific Return of Results Plans

Gene selection by sites for inclusion on the eMERGEseq panel was driven by both clinical and research needs leading to a final list for panel design of 109 genes, including the “ACMG56”^2^ and 53 additional site selected genes. Evidence review using the ClinGen gene-disease validity framework identified 35 of the additional 53 genes as having definite or strong association to disease. These genes were considered for further actionability analyses (See Figure 3). Most of the 18 genes with lower levels of validity were included by sites to enable research on these genes, reflecting the diverse goals of the eMERGE network including discovery as well as return of results.

**Figure 3:**
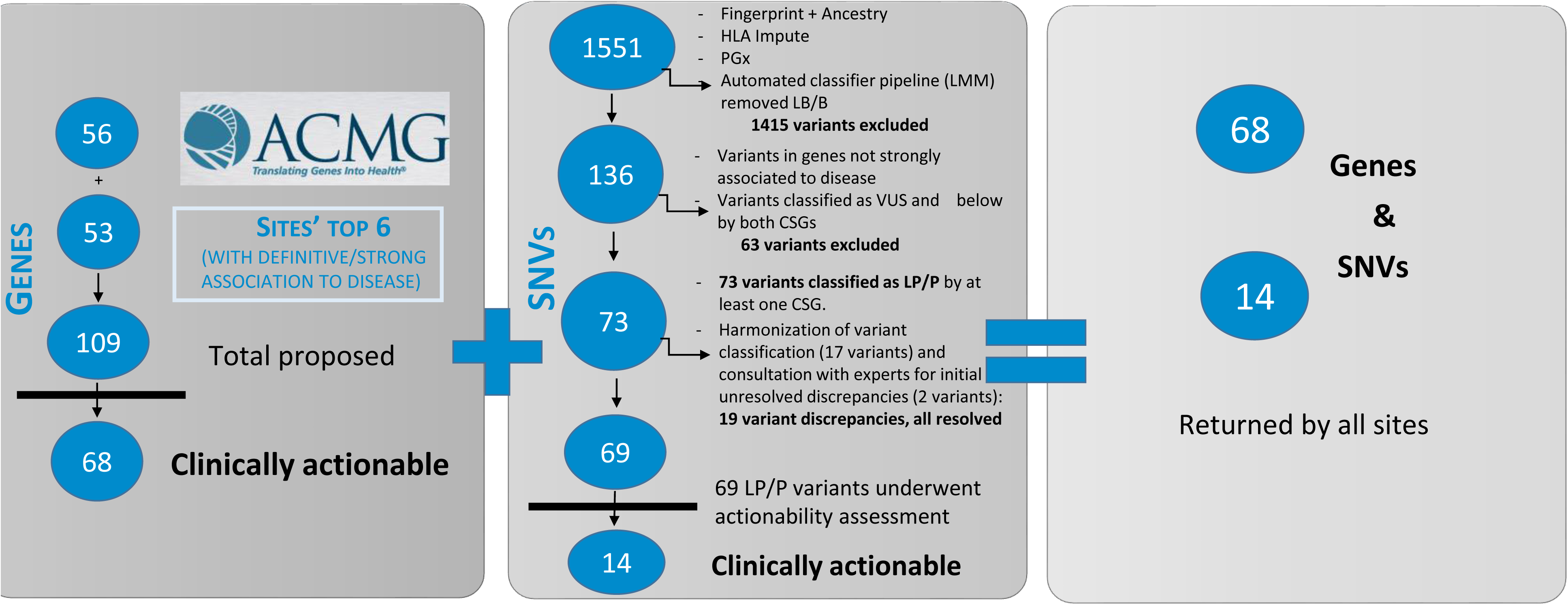
Content Development for the eMERGEseq Panel. Left panel: ClinGen gene-disease validity assessment for all site top 6 proposed genes. Those with definite and strong association to disease were considered for further actionability analyses. Middle panel: Clinical assessment for a subset of single nucleotide variants (SNVs). Those deemed Pathogenic/Likely Pathogenic were considered for actionability analyses. Right panel: Final consensus list of returnable content. This included all the ACMG56 genes, in addition to 12 genes and 14 variants were deemed actionable by the eMERGE Clinical Annotation Working Group.

A subset of the 1415 site submitted SNVs were for fingerprinting and ancestry, HLA, or PGx categories, or have been previously classified as likely benign or benign and were thus excluded for further analyses of potential pathogenicity. The remaining 136 variants were considered for further clinical assessment. Seventy three variants were classified as either likely pathogenic or pathogenic by at least one of the SCs. Of these, 19 had discrepant classifications between the two SCs. These were resolved by variant re-assessment and scoring on published evidence as well as combined internal evidence from both SCs. For two variants, the eMERGE Clinical Annotation WG was consulted to assist in resolving interpretation differences. A final list of 69 pathogenic/likely pathogenic variants was established and further considered for actionability analyses (Figure 3).

The eMERGE Clinical Annotation WG evaluated the 35 non-ACMG56 strong/definitive genes and 69 associated pathogenic/likely pathogenic variants, based on whether there was a substantially increased risk of serious disease that could be prevented or managed differently if the risk were known. In addition to the ACMG56, 12 genes and 14 variants were deemed actionable by the eMERGE Clinical Annotation WG and placed on a consensus list of returnable content (Table 2, Figure 3). While sites agreed that this list represented content that would generally be returnable, some sites requested modifications be made to the consensus list based to their return of results plans (Figure 4). For example, of the 11 sites, one that included pediatric biobank participants opted not to report variants in genes that increase risk of adult onset diseases but are not actionable during childhood. Also, not all sites chose to return *HFE* p.Cys282Tyr homozygotes. Four other sites requested additional genes and SNVs that were not on the consensus list. A full list of the content that was returned for each site can be found in Table S3. Additionally, one site is returning variants of uncertain significance in 13 colorectal cancer genes for a subset of their samples derived from a colorectal cancer cohort. Another clinical site requested genotypes at twelve SNP sites associated with low-density lipoprotein cholesterol (LDL-C) risk be included on their report.

**Figure 4:**
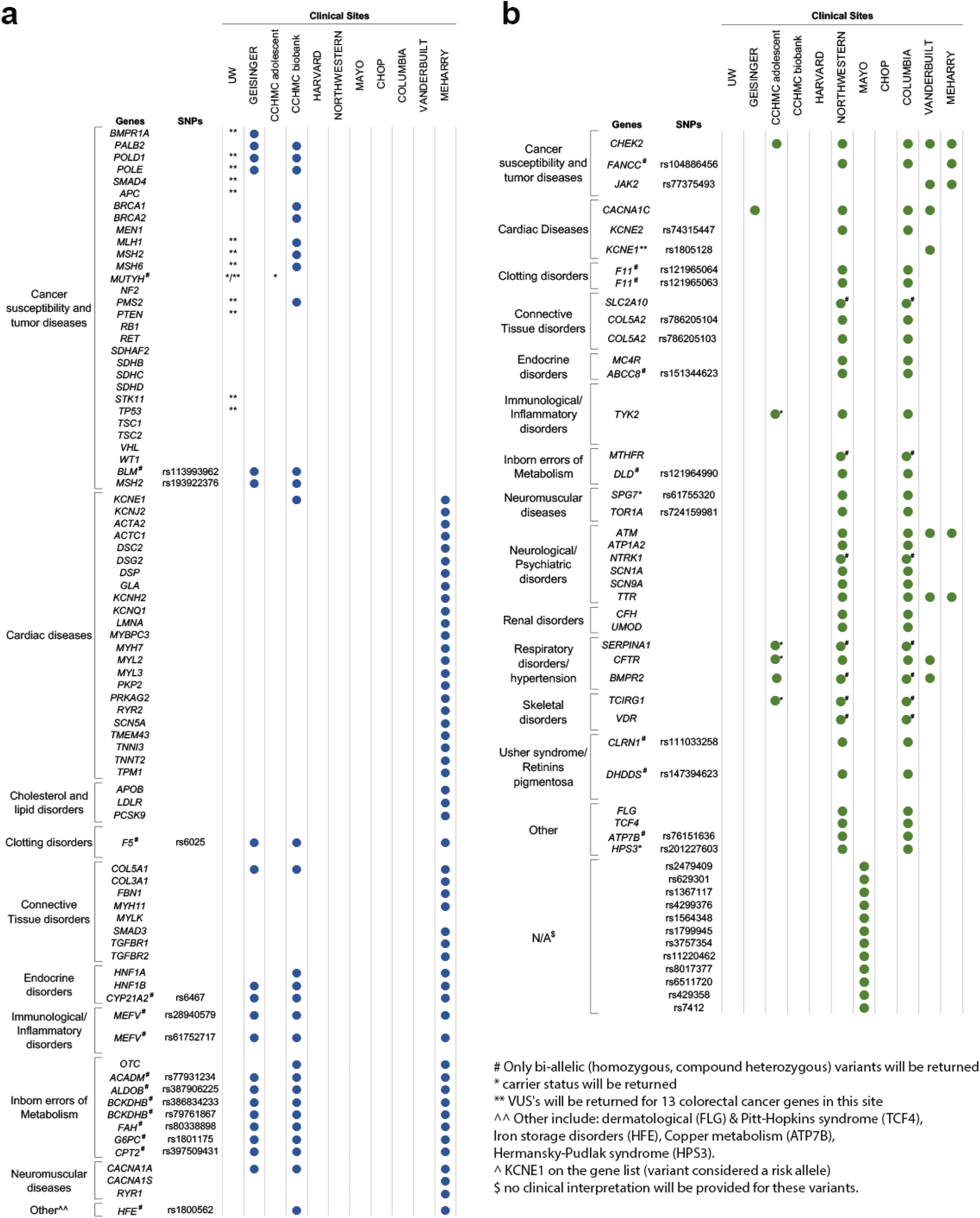
Site specific reportable list of genes/SNPs for which Pathogenic or Likely Pathogenic variants will be returned. a. Consensus List. Consensus list of returnable SNPs/Genes with site-specific exclusions indicated with a blue dot b. Site specific List. Non-consensus Genes/SNPs with site-specific inclusions indicated with a green dot

### (iv) Data Intake and Delivery

Data intake and delivery represented challenges for the network due to the plan to test distributed, heterogeneous EHR systems and other data sources used by sites and the need to deliver updated data interpretations. All demands were required to be met while managing issues of compliance and security for PHI protection. These challenges mimicked real-world situations as these are identical needs for any health care organization opting to interact with a research enterprise or reference laboratory. The data management required the development of three main informatic components:

#### (a) Data intake

Data intake and accessioning for each site was facilitated by an agreement of the specific PHI metadata to be supplied with each sample, as well as an agreement of a set of required ‘indications for testing’ that represented the primary phenotype data that tracked each sample through the network (see Methods).

#### (b) Clinical reporting

Within each pipeline, the standard validated product was a pdf report that was returned to the clinical investigators (see supplementary material for examples of reports: (Sample eMERGE report HGSC-CL, Sample eMERGE report HGSC-CL XML, Sample eMERGE report Partners Broad, Sample eMERGE report Partners Broad XML). Each clinical site had custom requirements for the report content, that reflected local preferences for data to be returned to patients. Each SC also had different reporting requirements – for example, some sites requested negative reports, others only returned positive reports^16^. Most sites also requested data in structured formats to enable direct integration onto their local EHRs.

The five clinical sites served by the Partners-Broad CSG received results delivered through the GeneInsight platform, which enabled storage and query of clinical reports. The six sites served by the BCM-HGSC utilized custom applications developed for report delivery. Possible difficulties in data sharing between different parts of the network were anticipated and obviated by development of an agreed .xml standard. This standard was based upon the GeneInsight specifications and facilitated communication across all components (See Methods and^17^).

The clinical sites therefore had two options –they could either use a stand-alone tool for report data management or alternatively the report data could be parsed into local customized systems.

For PGx data, in addition to receiving results in pdf reports (either individual reports by the BCM-HGSC, or batch reports by Partners-Broad), a standardized data format was also developed to deliver structured PGx data in the form of both variant level and diplotype results allowing sites to directly integrate PGx results into the EHR for clinical decision support.

#### (c) Research and Discovery via Data Commons and the De-identified Case Repository

Finally, the network required all deidentified data to reside together, to enable data mining for both basic research and to better inform clinical decision making with access to larger clinical datasets. There were two independent but complementary mechanisms for this. First, the GeneInsight tool maintains a record of all returned variant data from both sites in a de-identified case repository allowing an easy search interface for clinically reported variants.

A second site maintained the full set of eMERGE raw data in a cloud environment, managed by a middle-ware vendor, DNAnexus. This ‘eMERGE Commons’ was structured to house each DNA sequence file in the BAM format, as well as the annotations for the data in a vcf format. As clinical report delivery for the data generated in the Baylor SC also utilized the DNAnexus infrastructure, the full set of identified clinical reports and de-identified raw data were both resident in the cloud. The access permissions for the data were managed to allow only the clinical providers to access their patients’ clinical reports. The full set of raw data was available to all eMERGE investigators as sensitive PHI information had been removed.

### (v) Variant Interpretation Harmonization

To ensure consistency of results being returned across the eMERGE consortium, variant interpretation was harmonized between the SCs (Figure 5). In a pre-test launch, both SCs exchanged variants in reportable genes from their respective databases, totalling 23,663 unique variants. Of those, 1047 were previously classified by both SCs. The pre-test lauch data exchange showed 90% concordance in variant classification among variants classified as VUS, likely pathogenic and pathogenic by at least one SC. When likely pathogenic and pathogenic variants were grouped together, the concordance was 93%. When all variant classifications were considered, including benign vs likely benign, the data showed a 67.5% concordance. However, only 28, or 3% of the variants were deemed to affect reporting (VUS vs pathogenic 1.9%, VUS vs likely pathogenic 1.1%). The two SCs resolved all differences that would affect inclusion on clinical reports (i.e. pathogenic/likely pathogenic versus VUS).

**Figure 5:**
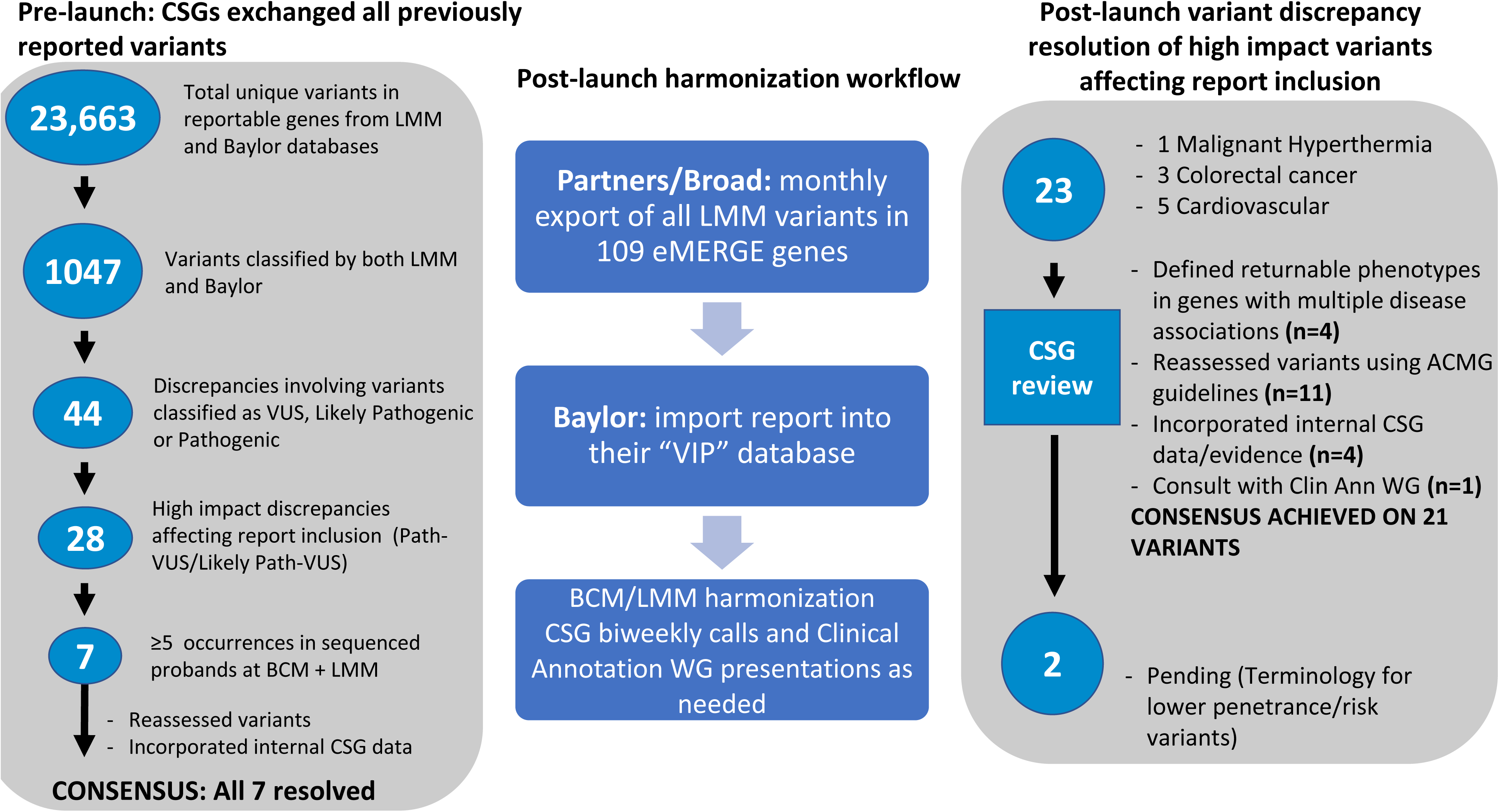
Variant harmonization process overview. Left panel: Pre-launch and post-launch harmonization processes involving the exchange of variants in reportable genes between the sequencing centers and the identification, prioritization and the resolution of discrepancies affecting report inclusion.

An ongoing process was also developed to ensure continuous harmonization of variant interpretation (Figure 5). As of October 2018, 23 initial discrepancies of interpretation of variants from five disease areas were considered, based upon potential to affect report inclusion. Most variants (83%) were immediately resolved when reassessed by the SCs, using ACMG guidelines, incorporating additional laboratory-specific evidence, after defining returnable phenotypes in genes with multiple disease associations (for example malignant hyperthermia vs. myopathy for *RYR1*), or defining terminology for lower penetrance/risk variants. For one variant, resolution required input from additional eMERGE investigators through the eMERGE Clinical Annotation WG.

Three variants (p.Ile1307Lys in *APC*, p.Met54Thr in *KCNE2*, and p.Asp85Asn in *KCNE1*) were noteworthy as the interpretations were more discrepant upon initial assessment (i.e. ‘two-steps’: pathogenic vs likely benign), although the evidence used by both centers was identical. These represented variants that have significantly reduced penetrance, leading to difficulties applying the ACMG/AMP classification framework, which is designed primarily for highly penetrant Mendelian disorders. Nevertheless, some sites chose to return the *APC* variant as it imparts a two-fold risk of colorectal cancer in Ashkenazi Jewish individuals, even though its effect in other populations in unclear. Other sites elected to return the *KCNE2* variant, as it has been associated with variable presentations such as arrhythmias, sinus bradycardia and long QT syndrome^18-201^. This type of classification discordance highlights the need for guidance on classification terminology for low penetrance variants for not only the eMERGE network, but for the entire medical genetics community.

### (vi) Return of Results and Aggregate Findings

As of December 2017, 15,754 cases had been collected and analyzed via the eMERGEseq panel. To coordinate analyses, these samples were included in a ‘data freeze’ termed ‘eMERGEseq Data Freeze 1.0’ (see supplementary methods). All of the 15,754 data freeze samples have passed through at least the primary variant assessment and review stage. For these assessed cases, a total of 1,913,377 variants were detected. A subset of these were excluded from further analyses due to a LB/B classification by the SCs or by an auto-classification pipeline based on allele frequency thresholds, or for having a low quality score. The remaining variants underwent a filtration process which returns a) predicted loss of function variants with a minor allele frequency (MAF) <1%, b) variants previously classified by the SCs as Likely Pathogenic(LP)/Pathogenic(P) regardless of MAF, and c) ClinVar P/LP as well as HGMD "DM” variants with a MAF<5%. This pipeline resulted in 4786 unique variants requiring further assessment. After expert review, these were further categorized as Benign (1%), Likely Benign (8%), VUS (69%), LP (7%), P (12%) or deemed as low penetrance risk alleles (0.5%). In addition, 95 unique copy number variants have been detected across the reviewed samples, with 74 gains and 34 losses. Of these, 35% were deemed reportable and were returned to sites. In summary, these data lead to a total of 679 cases projected to have a LP/P variant that would require a positive report to be issued.

Results being returned to sites currently fall into three categories: 1) Indication-based returnable results that include all sequence and copy number variants related to the site-provided indication for testing, 2) nonindication-based consensus returnable results that include all sequence and copy number variants in genes and SNVs comprising the consensus list of returnable content (see clinical content validation section) that are not related to indication for testing, and thus considered secondary findings and 3) non indication-based site-specific returnable results which include variants in additional site-requested genes that are not on the consensus list and not related to the indication for testing. Additionally, both SCs are returning results on pre-selected PGx SNVs as either addendums to individual patient reports or in a batch report that contains up to ∽185 samples (See Methods).

The positive rate for each category of findings is depicted in Figure 6. For the 15,754 cases that have been reviewed, 5,909 (38%) had an indication for testing. Of these, 115 (1.95%) had positive findings relevant to the indication for testing. Moreover, of the 15,754 individuals sequenced, 681 (4.5%) had additional/secondary findings of medical significance in genes and SNVs from the consensus list, that are being returned to participants. About 4,073 participants (26%) were enrolled in sites who were interested in returning Pathogenic and/or Likely Pathogenic variants in additional genes or SNVs that were not on the consensus list. In 153 cases (3.5%), a non-indication based, site specific returnable Pathogenic or Likely Pathogenic variant was identified. About half of these variants were in the *CHEK2* tumor suppressor gene, and are associated with an increased risk for a variety of cancers. Other variants were found in genes associated with cardiac disease, familial hypercholesterolemia and hemochromatosis (Figure 6). For indication-based assessments, detection rates were highest for hyperlipidemia (44%), colorectal cancer/polyps (34%) and breast/ovarian cancer (19%). Some phenotypes had no disease-causing variants identified due to either the absence of genes causative for the disorders on the eMERGEseq panel or the lack of a clear monogenic disease etiology for the disorder (e.g. abnormality of pain sensation, pediatric migraine). The rate of Pathogenic/Likely Pathogenic variants detected in participants without a clinical indication differed from site to site, ranging from 1.8% to 17%, depending upon the basis for participant selection, which were reflective of the underlying study designs of the individual sites. The overall positive rate for secondary findings was skewed higher for one site given that 1251 participants of the Geisinger cohort were preselected for a suspicious variant in a parallel exome study^21^. On the other hand, 2 sites had lower rates than expected either because their cohort had an indication related to genes in the secondary findings list that led to the removal of these genes from secondary findings reporting or because the site did not choose to return all results from the consensus list. When data from Geisinger participants preselected for a suspicious variants were excluded, the frequency of secondary findings was similar across sites, ranging from 2.7% to 4.9%, suggesting that the complexity of the network did not distort these results, and reflecting the success of the data and process harmonization. A further analysis of the factors that influence the rate of secondary findings return is underway (A. Gordon et al., 2018, American Society of Human Genetics, abstract).

**Figure 6:**
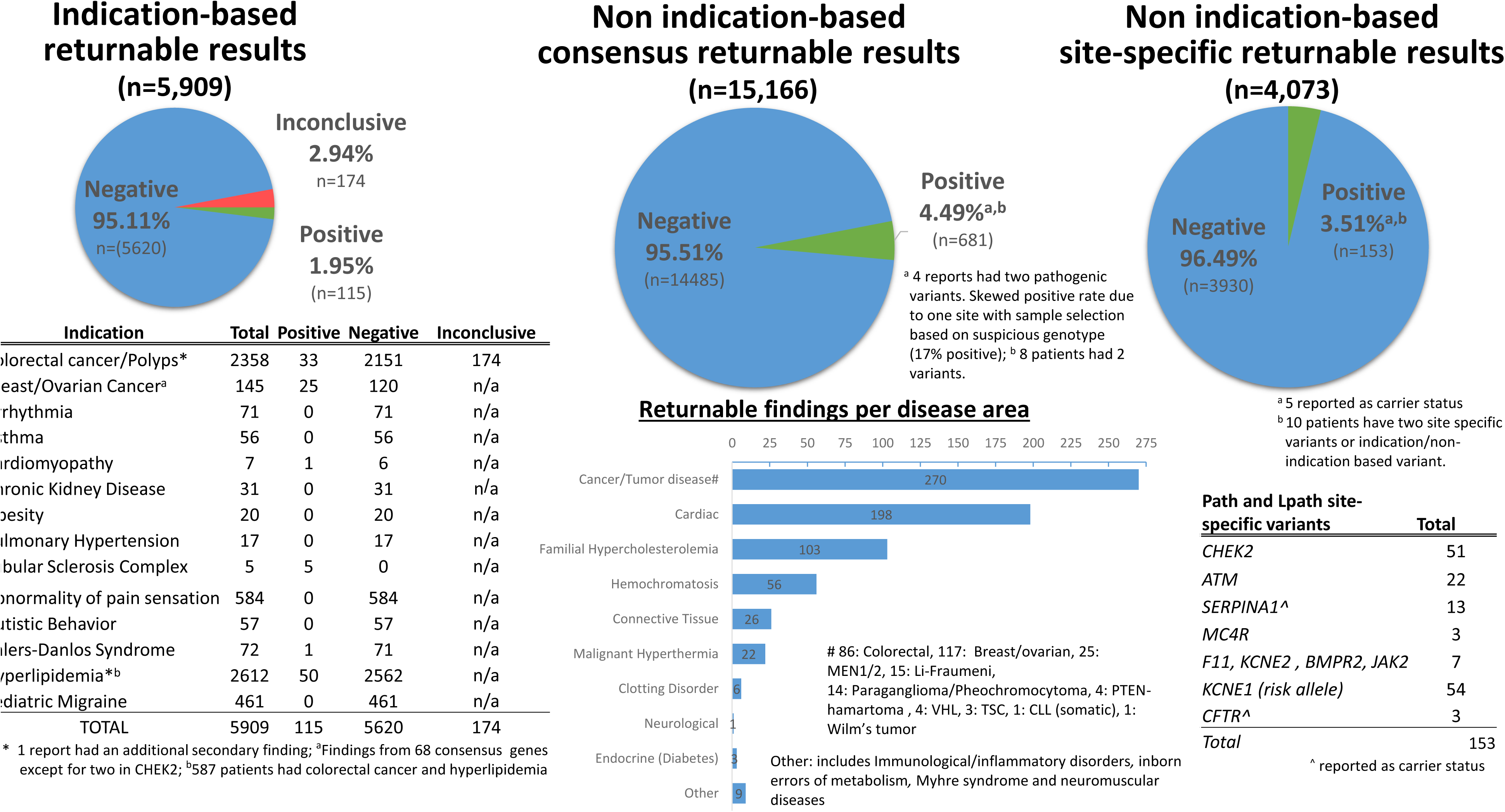
Aggregate findings returned to sites. The positive rate for each category of returnable findings for 15,754 participants from the eMERGESeq Data Freeze 1.0 is shown. For those with an indication for testing, the different indications are depicted (left). Secondary findings from the consensus gene list across the entire Data Freeze 1.0 cohort are broken down per disease area (middle). For a subset of participants, the number of Pathogenic and Likely Pathogenic variants in site-specific additional genes that are not on the consensus list are shown (right).

For PGx results, reports depicting genotype and related diplotype data, including whether the reported diplotype for each gene and resulting phenotype would result in a recommendation to modify dosage, have been used for approximately 9,000 participants from seven sites. Overall, the frequency of the reported diplotypes were concordant with the CPIC published frequency tables for each major race/ethnic group^22^.

One difference for diplotype interpretation was particularly informative. When both rs1800460 and rs1142345 are identified in the thiopurine methyltransferase *(TPMT*) gene, it cannot be ascertained whether these variants are in *cis*, resulting in a *TPMT*1/*3A* diplotype and intermediate metabolizer phenotype, or in *trans*, resulting in a *TPMT*3B/*3C* diplotype and a poor metabolizer phenotype. One SC emphasized the more common diplotype in their report, while the other emphasized the higher risk of the rarer diplotype under some drug regimens. With input from the sites and the eMERGE PGx working group, it was decided that the more common genotype would be reported with a warning that the rarer genotype could not be ruled out.

The majority of returned data reflected variants with relatively clear interpretations for participants, with variants that either had a large body of published evidence or were straightforward to interpret. Several cases, however, reflect interesting and unexpected findings.

The first finding involved what appeared to be a whole chromosome gain of chromosome 12. An NGS-based CNV calling algorithm detected a gain in all exons of six eMERGEseq genes on chromosome 12 *(CACNA1C, PKP2, VDR, MYL2, HNF1A* and *POLE*), which was confirmed by ddPCR. The *CACNA1C* and *POLE* genes are located near the telomeric end of the chromosome 12 p- and q-arms respectively, supporting a whole chromosome gain. Given that chromosome 12 trisomies are embryonic lethal, this CNV was assumed to be either of somatic origin or occuring as a mosaic variant. The former scenario is more likely as trisomy 12 is the most common somatic chromosomal aberration in chronic lymphocytic leukemia (CLL) but has also been observed in other B-cell lymphoproliferative disorders and is associated with a less favorable prognosis^23^. Rarely, trisomy 12 has been reported as a mosaic variant in individuals with a variety of clinical phenotypes ranging from reportedly normal to multiple congenital anomalies, dysmorphic features and developmental delay^24–28^. Most of these were identified prenatally, with less than 10 cases reported postnatally and even fewer detected in peripheral blood (for reviews see^27–28^). Additional clinical information provided by the site indicated that this patient has a complex medical history including diabetes, heart disease and a diagnosis of colorectal cancer at 87. While this finding is from a blood draw in early January 2016, this individual’s last complete blood count in 2010 showed no evidence of increased lymphocytes or any other abnormality suggesting a CLL diagnosis. While this type of result was not anticipated within the reporting scope for eMERGE III, upon further consultation with the site, this finding was included in the clinical report of the individual to encourage additional testing and/or management.

A second case with unexpected findings was associated with another copy number variant call. A duplication for all exons of the *OTC* and *GLA* genes, confirmed by ddPCR, was observed in a 40-year-old male not selected for phenotype. These genes are the only two present on the X chromosome on the eMERGEseq panel. Given that *OTC* and *GLA* are on the p and q arms respectively, the observed duplication is most likely a single event spanning the entire X chromosome. This is most consistent with a male with Klinefelter syndrome (47,XXY). Additional clinical information provided by the site confirmed a prior diagnosis of Klinefelter syndrome that had been confirmed by chromosomal karyotyping. Although a clinical report was not issued for this individual, these findings serve to further validate the sensitivity of NGS-based copy number calling.

The third unexpected category of findings was that six individuals presented with apparently mosaic variants in genes that predispose to cancer or cardiomyopathy *(TP53, CHEK2, ATM, MYH7).* The presence of mosaics was based upon the ascertainment of allelic variants that were present in <30% of the DNA sequence reads at the variant site. Initial observations were screened manually to eliminate false positives due to mis-mapping to pseudogene sites or other technical errors. The presence of the mosaic variants was subsequently confirmed by Sanger sequencing and clinical reporting offered to the referring sites.

## DISCUSSION

The introduction of clinical sequencing into the phase III of the eMERGE network has provided a framework for large-scale clinical translation of genomic data in healthcare, as well as for the seamless integration of research studies into clinical data management. The network integrated a large number of research groups with diverse interests, and a common mission to deliver genomic health care. To stimulate and address challenges for the delivery of genomic medicine, a large number of samples were tested and state of the art methods for interpretation and data delivery were applied.

A primary driver for the study design was cost and therefore a gene-panel was chosen as a primary platform for genomic analyses. Whole exome sequencing was considered. However, while exomes would have offered increased flexibility and saved time in design and testing, the network determined that a more focused target of ∽100 genes was needed to stay within the budget for testing all 25,000 participants. In addition, sites individually contributed research data on subjects using high density genotyping arrays allowing for genome-wide association studies which are not discussed here.

At the outset, the predictions were made as to the major challenges that would be faced and the most likely obstacles to achieving a smooth flow of clinical results, while maintaining access to research data. However, most of the challenges were not anticipated. For example, one challenge was the variety of different consents used to support the process as each site had a unique consent form and approach for their biobank and these consents sometimes stipulated requirements inconsistent with the network-wide decisions being made. As each site’s sequencing got started, these types of site-specific challenges were uncovered. Many sites altered their decisions around the reportable content and details of their reporting needs (e.g. which genes were reportable; whether negative reports were needed; whether reports should contain certain recommendations for genetic counseling, etc). There was evolving work around how to structure pharmacogenomic results to flow into EHRs and work to ensure the accurate provision of phenotypes from the sites to the SCs. One site needed accommodation for lower DNA input. These startup ‘hiccups’ led to significant delays in getting each site started with their sequencing and clinical reports. However, once a smooth workflow was developed for each site, the SCs were able to ramp up the rate of sequencing, interpretation and reporting. For example, during the first half of the project 5713 cases were completed, versus 9525 cases completed during the second half.

## CONCLUSIONS

An important outcome of the study is the generation of real data that reflects the practicality of such a large-scale biobank study. The network has provided an accurate estimate of the frequency of returnable results within the interrogated gene set. Further, the study has established the ability for two sequencing sites to adequately harmonize both the technical and interpretive aspects of clinical sequencing tests, a critical achievement to the standardization of genomic testing. Furthermore, the eMERGE network has accomplished the integration of structured genomic results directly into multiple electronic health record systems, setting the stage for the use of clinical decision support to enable genomic medicine.

### Consortia

The full list of members of the eMERGE consortium along with their affiliations and declarations of interests can be found in Table S5 (to be submitted in an upcoming revision).

Debbie Abrams, Samuel Adunyah, Majid Afshar, David Albers, Ladia Albertson-Junkans, Jen Albrecht, Darren Ames, Armand Antommaria, Paul Appelbaum, Krishna Aragam, Sandy Aronson, Sharon Aufox, Larry Babb, Adithya Balasubramanian, Shawn Banta, Melissa Basford, Joan Bathon, Christopher Bauer, Samantha Baxter, Meckenzie Behr, Barbara Benoit, Ashwini Bhat, Elizabeth Bhoj, Sue Bielinski, Sarah Bland, Paula Blasi, Carrie Blout, Eric Boerwinkle, Scott Bolesta, Kenneth Borthwick, Erwin Bottinger, Deb Bowen, Mark Bowser, Harrison Brand, Carmen Radecki Breitkopf, Murray Brilliant, Wendy Brodeur, Kevin Bruce, Adam Buchanan, Andrew Cagan, Pedro Caraballo, David Carey, David Carrell, Andrew Carroll, Robert Carroll, Peter Castaldi, Berta Almoguera Castillo, Lisa Castillo, Victor Castro, Bridget Chak, Gauthami Chandanavelli, Chia Yen Chen, Theodore Chiang, Rex Chisholm, Ken Christensen, Wendy Chung, Chris Chute, Brittany City, Ellen Clayton, Beth Cobb, Francis Cocjin, John Connolly, Nancy Cox, Paul Crane, Katherine Crew, David Crosslin, Damien Croteau-Chonka, Renata Pellegrino da Silva, Mariza De Andrade, Jessica De La Cruz, Emma Davenport, Colleen Davis, Dan Davis, Lea Davis, Matt Deardorff, Josh Denny, Shawn Denson, Tim Desmet, Parimala Devi, Keyue Ding, Michael Dinsmore, Sheila Dodge, Qunfeng Dong, Elizabeth Duffy, Phil Dunlea, Todd Edwards, Digna Velez Edwards, Mitchell Elkind, Christine Eng, Angelica Espinoza, Xiao Fan, Anna Farrell, David Fasel,

Alex Fedotov, Qiping Feng, Joseph Finkelstein, Mark Fleharty, Chamith Fonseka, Robyn Fossey, Andrea Foster, Robert Freimuth, Christopher Friedrich, Tanya Froehlich, Malia Fullerton, Lucinda Fulton, Birgit Funke, Stacey Gabriel, Vivian Gainer, Carlos Gallego, Nanibaa' Garrison, Tian Ge, Ali Gharavi, Richard Gibbs, Andrew Glazer, Joe Glessner, Jessica Goehringer, Adam Gordon, Chet Graham, Robert Green, Justin Gundelach, Jyoti Gupta, Hakon Hakonarson, Chris Hale, Taryn Hall, Maegan Harden, John Harley, Margaret Harr, Andrea Hartzler, Ali Hasnie, Geoff Hayes, Scott Hebbring, Nora Henrikson, Tim Herr, Andrew Hershey, Christin Hoell, Ingrid Holm, Kayla Howell, George Hripcsak, Alexander Hsieh, Jianhong Hu, John Hutton, Jodell Linder, Gail P. Jarvik, Joy Jayaseelan, Yunyun Jiang, Darren Johnson, Laney Jones, Sarah Jones, Yoonie Joo, Sheethal Jose, Navya Shilpa Josyula, Hayan Jouni, Ann Justice, Sara Kalla, Divya Kalra, Elizabeth Karlson, Sekar Kathiresan, Dave Kaufman, Kenneth Kaufman, Melissa Kelly, Eimear Kenny, Dustin Key, Abel Kho, Les Kirchner, Krzysztof Kiryluk, Terrie Kitchner, Barbara Klanderman, Derek Klarin, Eric Klee, Rachel Knevel, Dennis Ko, David Kochan, Barbara Koenig, Viktoriya Korchina, Leah Kottyan, Christie Kovar, Joel Krier, Emily Kudalkar, Rita Kukafka, Erin Kulick, Iftikhar Kullo, Philip Lammers, Eric Larson, Jennifer Layden, Joe Leader, Matthew Lebo, Magalie Leduc, Mike Lee, Yvonne Lee, Niall Lennon, Kathy Leppig, Nancy D. Leslie, Bruce Levy, Matthew Lewis, Dingcheng Li, Rongling Li, Wayne Liang, Chiao-Feng Lin, Noralane Lindor, Todd Lingren, James Linneman, Hongfang Liu, Wen Liu, Xiuping Liu, Yuan Luo, John Lynch, Hayley Lyon, Daniel Macarthur, Alyssa Macbeth, Harshad Mahadeshwar, Lisa Mahanta, Brad Malin, Vishnu Mallipeddi, Teri Manolio, Brandy Mapes, Maddalena Marasa, Talar Markossian, Keith Marsolo, Jen McCormick, Michelle McGowan, Elizabeth McNally, Ana Mejia, Jim Meldrim, Kelly Melissa, Frank Mentch, Jonathan Mosley, Shubhabrata Mukherjee, Tom Mullen, Jesus Muniz, David Raul Murdock, Shawn Murphy, John Murray, Mullai Murugan, Donna Muzny, Melanie F. Myers, Bahram Namjou, Pradeep Natarajan, Yizhao Ni, William Nichols, Nahal Nikroo, Laurie Novack, Aniwaa Owusu Obeng, Adelaide Arruda Olson, Janet Olson, Robb Onofrio, Casey Overby-Taylor, Jen Pacheco, Brian Palazzo, Melody Palmer, Jyoti Pathak, Peggy Peissig, Sarah Pendergrass, Kelly Perry, Nate Person, Josh F. Peterson, Lynn Petukhova, Sandie Pisieczko, Kelly Pittman, Siddharth Pratap, Cynthia A. Prows, Rebecca Pulk, Alanna Kulchak Rahm, Ratika Raj, James Ralston, Arvind Ramaprasan, Andrea Ramirez, Luke Rasmussen, Laura Rasmussen-Torvik, Hila Milo Rasouly, Soumya Raychaudhuri, Heidi Rehm, Marylyn Ritchie, Catherine Rives, Beenish Riza, Jamie R. Robinson, Dan Roden, Elisabeth Rosenthal, Jason Ross, Megan Roy-Puckelwartz, Janey Russell,Mert Sabuncu, Senthikumar Sadhasivam, Mayya Safarova, William Salerno, Saskia Sanderson, Simone Sanna-Cherchi, Avni Santani, Dan Schaid, Steven Scherer, Cloann Schultz, Rachel Schwiter, Stuart Scott, Aaron Scrol, Soumitra Sengupta, Nilay Shah, Catherine Shain, Ning 'Sunny' Shang, Himanshu Sharma, Richard Sharp, Yufeng Shen, Ben Moore Shoemaker, Patrick Sleiman, Kara Slowik, Josh Smit, Maureen Smith, Jordan Smoller, Sara Snipes, Susan Snyder, Jung Son, Peter Speltz, Justin Starren, Paul Steele, Kimberly Strauss, Mary Stroud, Amy Sturm, Jessica Su, Agnes Sundaresan, Michael Talkowski, Peter Tarczy-Hornoch, Stephen Thibodeau, Will Thompson, Lifeng Tian, Kasia Tolwinski, Maurine Tong, Sue Trinidad, Meghna Trivedi, Ellen Tsai, Sara L. Van Driest, Sean Vargas, Matthew Varugheese, David Vawdrey, Lyam Vazquez, David Veenstra, Eric Venner, Miguel Verbitsky, Gina Vicente, Sander Vinks, Carolyn Vitek, Michael Wagner, Kimberly Walker, Nephi Walton, Theresa Walunas, Bridget Wang, Kai Wang, Liuyang Wang, Llwen Wang, Qiaoyan Wang, Julia Wattacheril, Firas Wehbe, Wei-Qi Wei, Scott Weiss, Georgia L. Wiesner, Quinn Wells, Chunhua Weng, Peter White, Cathy Wicklund, Ken Wiley, Janet Williams, Marc S. Williams, Mike Wilson, Robert Winchester, Erin Winkler, Leora Witkowski, Betty Woolf, Eric Wright, Tsung-Jung Wu, Julia Wynn, Yaping Yang, Zi (Carol) Ye, Meliha Yetisgen, Zachary Yoneda, Ge Zhang, Kejian Zhang, Lan Zhang, Hana Zouk.

## Authors’ contributions

### Leadership

Hana Zouk*, Eric Venner*, Donna Muzny, Niall Lennon, Heidi L. Rehm^#^, Richard A. Gibbs^#^

### Test Design, Validation, Metric Tracking, CNV detection

Niall Lennon*, Kimberly Walker*, Donna Muzny, Adam Gordon, Mark Bowser, Maegan Harden, Theodore Chiang, Elizabeth Duffy, Jianhong Hu, Matthew Lebo, Alyssa Macbeth, Lisa Mahanta, Eric Venner, Tsung-Jung Wu, Gail P. Jarvik, Hana Zouk, Heidi Rehm, Richard A. Gibbs, Birgit Funke^#^, Donna Muzny^#^

### Validity and Actionability Assessment for Genes and SNPs

Hana Zouk, Magalie Leduc, Emily Kudalkar, Adam Gordon, Clinical Annotation WG, Eric Venner, Heidi Rehm, Gail P. Jarvik, Birgit Funke

### Data Intake and Delivery

Larry Babb, Mullai Murugan, EHRI WG, Darren Ames, Will Salerno, Maegan Harden, Chet Graham, Lisa Mahanta, Matthew Lebo, Heidi Rehm, Richard A. Gibbs, Sandy Aronson, Eric Venner

### Pharmacogenomics Reporting

Barbara Klanderman, Hana Zouk, Eric Venner, Elizabeth Duffy, Chiao-Feng Lin, Chet Graham, Lisa Mahanta, Donna Muzny, Rebecca Pulk, Steven Scherer, Matthew Lebo, Richard A. Gibbs, Birgit Funke, Magalie Leduc

### Variant Interpretation Harmonization and Result Interpretation

Yunyun Jiang*, Leora Witkowski*, Yaping Yang, Christine Eng, Matthew Varugheese, Eric Venner, Clinical Annotation WG, Birgit Funke, Richard A. Gibbs, Heidi Rehm, Magalie Leduc^#^, Hana Zouk^#^

### Clinical Site Representatives for Return of Results

Armand Antommaria (CCHMC), Nancy D. Leslie (CCHMC), Melanie F. Myers (CCHMC), Cynthia A. Prows (CCHMC), Wendy Chung (Columbia), David Fasel (Columbia), Hila Rasouly Milo (Columbia), Chunhua Weng (Columbia), Scott Bolesta (Geisinger), Melissa Kelly (Geisinger), Rebecca Pulk (Geisinger), Marc S. Williams (Geisinger), Gail P. Jarvik (UW), Eric Larson (KPW), Kathleen Leppig (KPW), James Ralston (KPW), David Kochan (Mayo), Iftikhar Kullo (Mayo), Noralene Lindor (Mayo), Erin Winkler (Mayo), Christin Hoell (Northwestern), Laura J. Rasmussen-Torvik (Northwestern), Maureen E. Smith (Northwestern), Robert C. Green (Partners), Jordan W. Smoller (Partners), Josh F.Peterson (VUMC), Jamie R. Robinson (VUMC), Ben Shoemaker (VUMC), Sara L. Van Driest (VUMC), Quinn Wells (VUMC), Georgia L. Wiesner (VUMC).

### BCM-Human Genome Sequencing Center

Donna Muzny, Eric Venner, Jianhong Hu, Kimberly Walker, Sara Kalla, Theodore Chiang, Tsung-Jung Wu, Ritika Raj, Andrea Foster, Adithya Balasubramanian, Jesus Muniz, Shawn Denson, Gauthami Chandanavelli, Wen Liu, Harshad Mahadeshwar, Kimberly Strauss, Sean Vargas, Lan Zhang, Xiuping Liu, Qiaoyan Wang, Joy Jayaseelan, Darren Ames, Divya Kalra, Beenish Riza, Jessica De La Cruz, Brian Palazzo, LIwen Wang, William Salerno, Viktoriya Korchina, Christie Kovar, Yunyun Jiang, Magalie Leduc, David Raul Murdock, Eric Boerwinkle, Victoria Yi, Yaping Yang, Mullai Murugan, Christine Eng, Richard A. Gibbs

### Partners Laboratory for Molecular Medicine/Broad Institute Sequencing Center

Hana Zouk, Maegan Harden, Leora Witkowski, Samuel Aronson, Larry Babb, Samantha Baxter, Mark Bowser, Wendy Brodeur, Sheila Dodge, Phil Dunlea, Christopher Friedrich, Tim Desmet, Michael Dinsmore, Mark Fleharty, Chet Graham, Elizabeth D. Hynes, Emily Kudalkar, Matthew Lebo, Chiao-Feng Lin, Hayley Lyon, Alyssa Macbeth, Lisa Mahanta, Jim Meldrim, Tom Mullen, Robb Onofrio, Kara Slowik, Gina Vicente, Mike Wilson, Betty Woolf, Stacey Gabriel, Birgit Funke^#^, Niall Lennon^#^, Heidi L. Rehm^#^

### Coordinating Center

Melissa Basford, David Crosslin, Adam Gordon, Kayla Howell, Gail P. Jarvik, Jodell Linder (Jackson), Ian Stanaway, Josh Peterson (PI)

### eMERGE Working Group Co-Chairs

<u>Clinical Annotation:</u> Gail P. Jarvik and Heidi Rehm; <u>EHR Integration:</u> Sandy Aronson and Casey Overby-Taylor; <u>Genomics:</u> David Crosslin, Megan Roy-Puckelwartz and Patrick Sleiman; <u>Outcomes::</u> Hakon Hakonarson, Josh Peterson and Marc S. Williams; <u>PGx</u> Cynthia A. Prows and Laura Rasmussen-Torvik; <u>Phenotyping:</u> George Hripcsak and Peggy Peissig; <u>Return of Results/ELSI:</u> Ingrid Holm and Iftikhar Kullo.

### eMERGE Principal Investigators

Rex Chisholm (Steering Committee Chair), Samuel Adunyah, David Crosslin, Josh Denny, Ali Gharavi, Richard Gibbs, Hakon Hakonarson, John Harley, George Hripcsak, Gail P. Jarvik, Elizabeth Karlson, Iftikhar Kullo, Philip Lammers, Eric Larson, Niall Lennon, Shawn Murphy, Josh Peterson, Heidi Rehm, Dan Roden, Marylyn Ritchie, Richard Sharp, Maureen Smith, Jordan Smoller, Stephen Thibodeau, Chunhua Weng, Scott Weiss, Marc S. Williams

### NHGRI Staff

Rongling Li (Program Director), Ken Wiley (Program Director), Jyoti Dayal, Sheethal Jose, Teri Manolio (Division Director)

*^#^contributed equally

## DECLARATION OF INTERESTS

See Table S5 (to be submitted in an upcoming revision)

### Ethics approval and consent to participate

All 10 sample collection sites consented participants under Institutional Review Board-approved protocols and the two sequencing centers had IRB-approved protocols that deferred consent to the participating sites. Protocol numbers are as follows: Partners Healthcare (2015P000929), Baylor College of Medicine (#H-40455).

### Consent for publication

Not applicable

### Availability of data and material

The datasets generated and/or analysed during the current study will be publicly available in the dbGaP repository under phs001616.v1.p1 and pre-dbGaP submission access can also be requested on the eMERGE website https://emerge.mc.vanderbilt.edu/collaborate/

